# Colloidal aggregation confounds cell-based Covid-19 antiviral screens

**DOI:** 10.1101/2023.10.27.564435

**Authors:** Isabella S. Glenn, Lauren N. Hall, Mir M. Khalid, Melanie Ott, Brian K. Shoichet

## Abstract

Colloidal aggregation is one of the largest contributors to false-positives in early drug discovery and chemical biology. Much work has focused on its impact on pure-protein screens; here we consider aggregations role in cell-based infectivity assays in Covid-19 drug repurposing. We began by investigating the potential aggregation of 41 drug candidates reported as SARs-CoV-2 entry inhibitors. Of these, 17 formed colloidal-particles by dynamic light scattering and exhibited detergent-dependent enzyme inhibition. To evaluate antiviral efficacy of the drugs in cells we used spike pseudotyped lentivirus and pre-saturation of the colloids with BSA. The antiviral potency of the aggregators was diminished by at least 10-fold and often entirely eliminated in the presence of BSA, suggesting antiviral activity can be attributed to the non-specific nature of the colloids. In confocal microscopy, the aggregates induced fluorescent puncta of labeled spike protein, consistent with sequestration of the protein on the colloidal particles. Addition of either non-ionic detergent or of BSA disrupted these puncta. These observations suggest that colloidal aggregation is common among cell-based anti-viral drug repurposing, and perhaps cell-based assays more broadly, and offers rapid counter-screens to detect and eliminate these artifacts, allowing the community invest resources in compounds with true potential as a Covid-19 therapeutic.

## Introduction

An attractive approach to speeding early drug discovery is re-purposing approved drugs for new indications. Eliding hit identification, lead-optimization, lead-to-candidacy, extensive animal toxicology and much pharmacokinetics, drug repurposing can substantially cut the time and costs of drug development with apparently de-risked molecules [1, 2]. With the arrival Covid-19, there was great interest in this approach to rapidly advance drugs that could treat the virus [3, 4]. Accordingly, many screens of drug-library screens were undertaken, largely in academic and government laboratories. Early results of these screens suggested that there was a large and diverse group of approved drugs that had antiviral efficacy against the virus in cell-based and protein-based screens, at least at high nM to low µM concentrations of the drugs [5-7].

Several of the effective antiviral drugs to emerge in the pandemic did in fact reflect a form of drug repurposing. For instance, remdesivir was an antiviral repurposed from its original use against Ebola [8-10], Paxlovid had its origins in campaigns begun against SARS-1 [11, 12], while drugs like plitidepsin [13], and zotatifin [14], where clinical investigation is ongoing, were molecules with mechanisms of action known to have antiviral potential (e.g., inhibition of protein biosynthesis). There was a much larger class of molecules, however, that emerged from the screens without an apparent mechanism, but were phenotypically active in cell-based viral infectivity and replication assays [5, 15-19].

The lack of mechanism was a concern [20], and on mechanistic examination liabilities emerged. Among the drugs repurposed for cell-based antiviral activity, many were cationic amphiphiles that turned out to induce phospholipidosis and the disruption of lipid homeostasis, a toxicity-linked mechanism likely without useful anti-viral effect [21, 22]. A second class of repurposed drug was those found in pure protein inhibition studies. Many of these were subsequently found to act through colloidal aggregation [23]. Aggregation is a well-known artifact in early drug discovery [24-30], often acting in such pure protein assays, where the small molecules aggregate into colloidal particles that can be detected by dynamic light scatter (DLS), electron microscopy, and NMR spectroscopy [26, 31-33]. Particle size range from 50 nM to close to 1 micron [26, 34, 35], and in this form sequester, mildly denature [36], and inhibit the target protein with little apparent selectivity. While it may be surprising that “derisked” molecules like drugs will aggregate and artifactually inhibit proteins, at concentrations that are relevant for screening they are just as prone to do so as other molecules in screening decks [38, 39]. The phospholipidosis-inducing drugs and the colloidal aggregating drugs typically have little future as antivirals.

A third class among the repurposed drugs were those that were not cationic amphiphiles, and so unlikely to induce phospholipidosis, but were nevertheless active in cell-based screens [17-19, 40, 41]. While these assays did not involve the inhibition of pure proteins, which is where colloidal aggregation often has an impact, many nevertheless had physical properties that resemble those of known aggregators: hydrophobic, highly conjugated, and often with multiple phenols. These molecules are reported to act as entry inhibitors, preventing SARS-CoV-2 to infect cells, often via activity against the viral spike protein. Here we investigate the ability of colloidal aggregation to confound Covid-19 repurposing screens performed in cell-based assays. We focused on reported entry inhibitors, where protein sequestration by colloids seems plausible, testing 41 drugs for activity via an aggregation-based mechanism. Of these, 17 behaved as classic aggregators, forming particles by dynamic light scattering (DLS) and inhibiting counter-screening enzymes in the absence but not the presence of small amounts of non-ionic detergent. Another two did not form particles by DLS, but had the other harbingers of aggregation. For most these compounds, bovine serum albumin, which reverses aggregation-based inhibition in pure protein assays, also reversed inhibition of viral infectivity in pseudovirus assays. Taken together, these results support the idea that colloidal aggregation can also impact cell-based assays and in particular has led to false-positives among repurposed drugs for SARS-CoV-2. How this cell-based effect may be controlled for in future studies will be considered.

## Results

### Colloidal aggregators in SARS-CoV-2 antiviral cell-based screens

A literature search found 41 drug-library candidates to test for colloidal aggregation. Drug library molecules met the following criteria: they were reported entry inhibitors against SARS-CoV-2 or SARS in cell-based assays and were either previously reported as colloidal aggregators or structurally resembled known aggregators (e.g., conjugated ring systems and clogP typically ≥3; several were included that did not meet these criteria, mostly as internal controls) (**Table S1**) [17-19, 40-56]. We also tested one compound, amentoflavone that only had computational support for its role as an entry inhibitor, but had several reports of its potential as a COIVD-19 antiviral [57-61]. Among the 41 drug-library molecules screened, 17 met all of our criteria for colloidal aggregation. For a drug to be considered an aggregator it had to inhibit counter-screen enzymes AmpC β-lactamase (AmpC) or malate dehydrogenase (MDH) at relevant concentrations, inhibition had to be reversible by detergent or by BSA, and the molecule had to form particles by Dynamic Light Scattering (DLS). Often these molecules also had high Hill slopes, but this is not always true of colloidal aggregators [23, 27] and was not a strict criterion here.

We began by screening all compounds by DLS. If a compound formed colloid-like particles at a concentration of 100µM or 200µM (e.g., scattering intensity β1ξ10^6^ and particle size between 100 and 1000nm), we measured particle formation in concentration-response (**Figure 1)**. To measure critical aggregation concentrations (CACs), we separated non-aggregating and aggregating concentrations with a best fit line, as previously [23, 31, 62], with the point of intersection identifying the CAC for each drug library molecule (**Table 1**). Of the 41 molecules tested, 33 formed colloid-like particles by DLS (**Figure S1**), with CAC values ranging from 1 to 83 µM. Typically, these values fell within the range of the literature reported IC_50_ values for the entry inhibitors, and in seven cases with CACs below the reported IC_50_ values.

**Figure 1.**
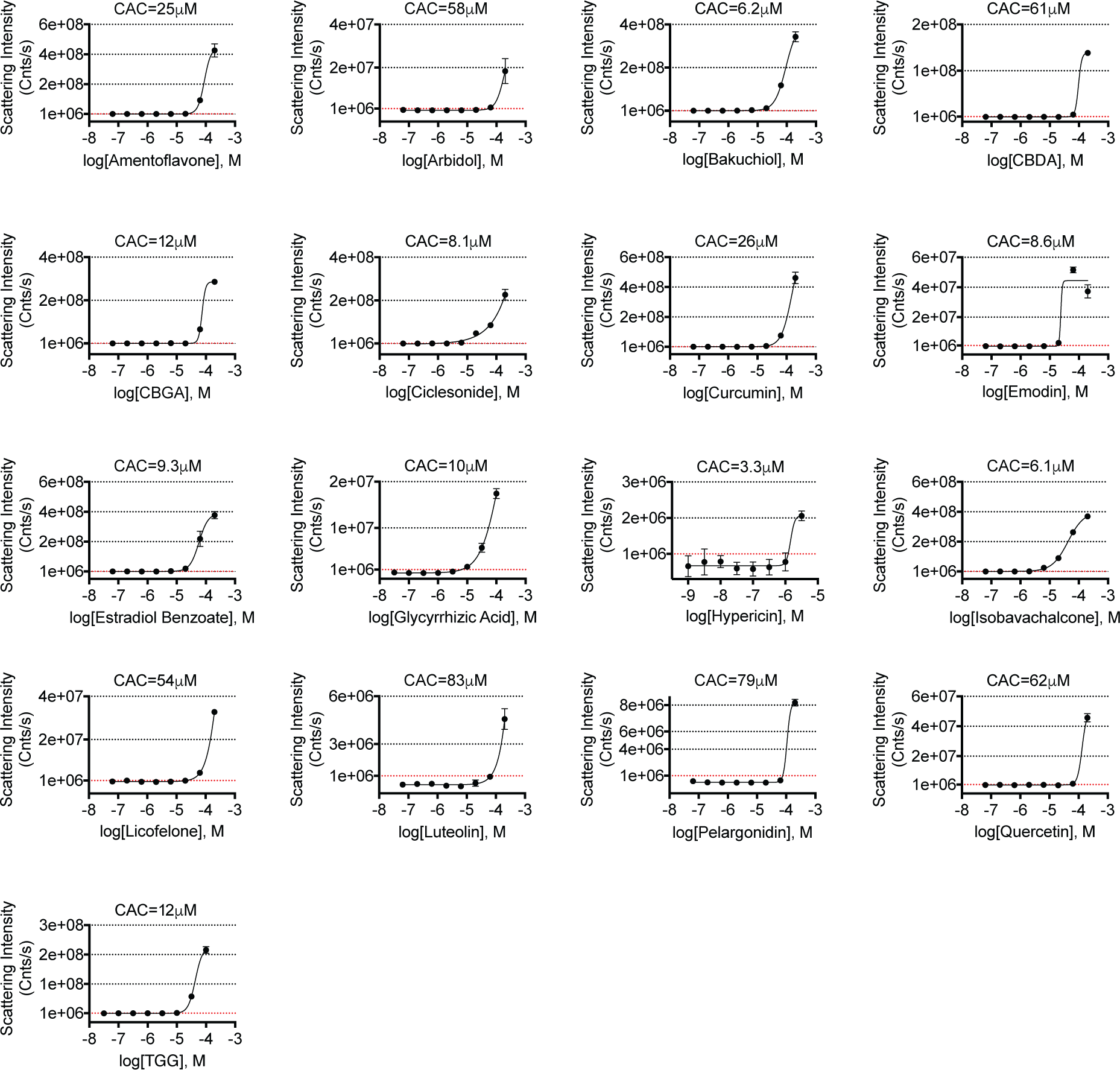
Dynamic light scattering performed on colloidal aggregator candidates by scattering intensity threshold of 1x10^6^. Point of intersection serves as CAC value for each compound. All measurements were performed in triplicate.

**Table 1.**
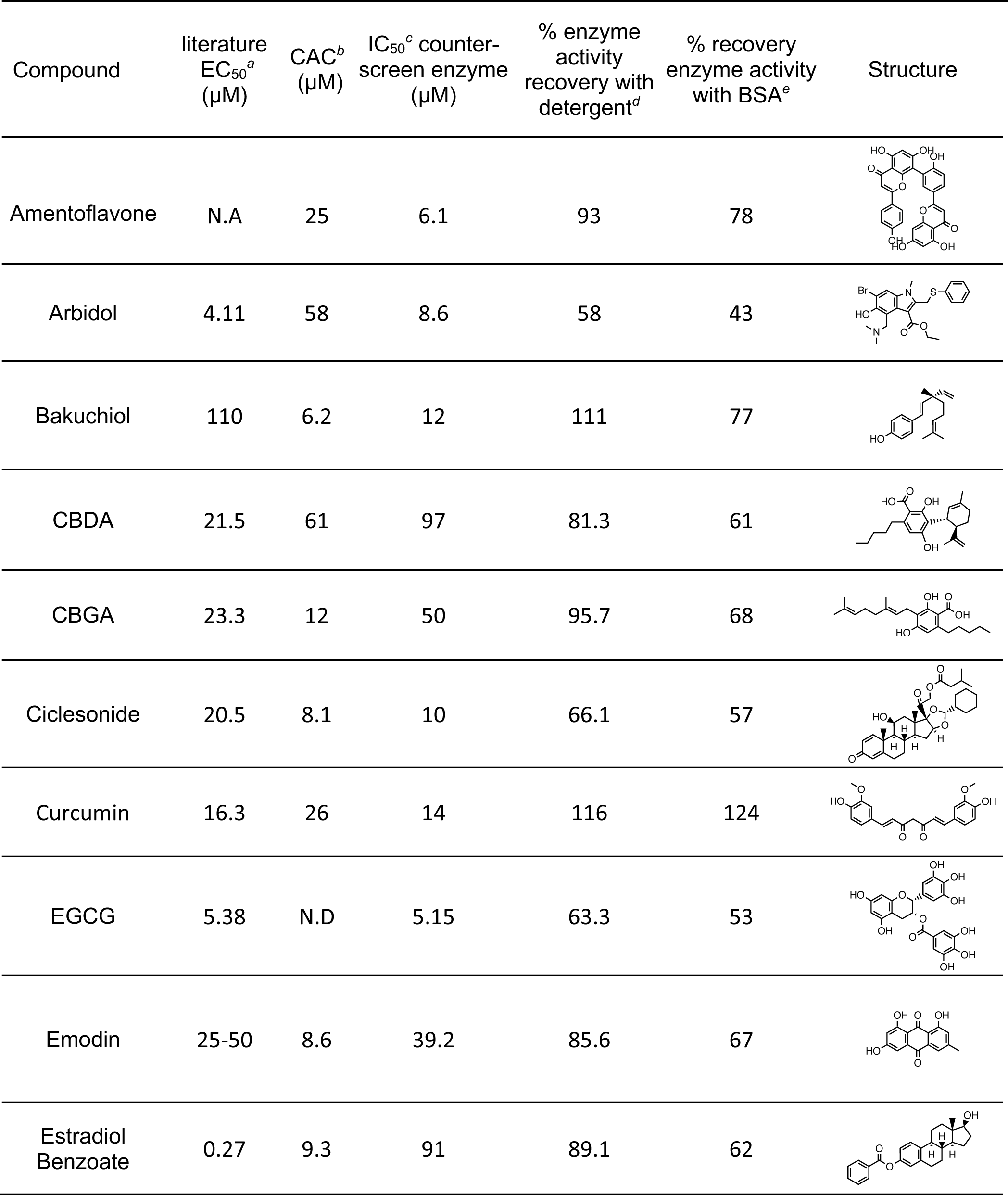
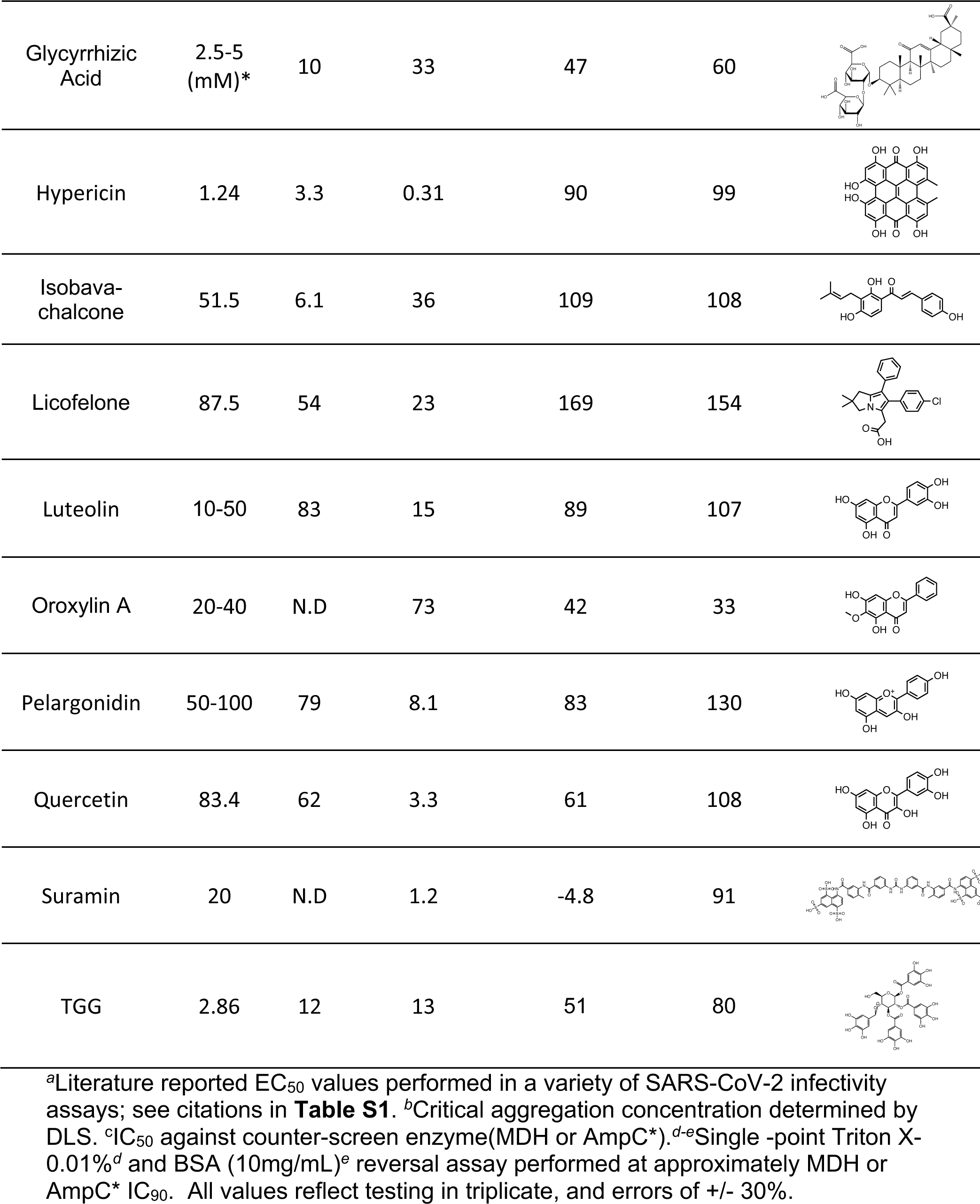
SARS-CoV-2 repurposed drugs that cause colloidal aggregation.

| Compound | literature<br>EC<sub>50</sub><sup>a</sup><br>(μM) | CAC<sup>b</sup><br>(μM) | IC<sub>50</sub><sup>c</sup> counter-screen enzyme<br>(μM) | % enzyme<br>activity<br>recovery with<br>detergent<sup>d</sup> | % recovery<br>enzyme activity<br>with BSA<sup>e</sup> | Structure |
| --- | --- | --- | --- | --- | --- | --- |
| Amentoflavone | N.A | 25 | 6.1 | 93 | 78 |  |
| Arbidol | 4.11 | 58 | 8.6 | 58 | 43 |  |
| Bakuchiol | 110 | 6.2 | 12 | 111 | 77 |  |
| CBDA | 21.5 | 61 | 97 | 81.3 | 61 |  |
| CBGA | 23.3 | 12 | 50 | 95.7 | 68 |  |
| Ciclesonide | 20.5 | 8.1 | 10 | 66.1 | 57 |  |
| Curcumin | 16.3 | 26 | 14 | 116 | 124 |  |
| EGCG | 5.38 | N.D | 5.15 | 63.3 | 53 |  |
| Emodin | 25-50 | 8.6 | 39.2 | 85.6 | 67 |  |
| Estradiol Benzoate | 0.27 | 9.3 | 91 | 89.1 | 62 |  |
| Glycyrrhizic Acid | 2.5-5 (mM)* | 10 | 33 | 47 | 60 |  |
| Hypericin | 1.24 | 3.3 | 0.31 | 90 | 99 |  |
| Isobavachalcone | 51.5 | 6.1 | 36 | 109 | 108 |  |
| Licofelone | 87.5 | 54 | 23 | 169 | 154 |  |
| Luteolin | 10-50 | 83 | 15 | 89 | 107 |  |
| Oroxylin A | 20-40 | N.D | 73 | 42 | 33 |  |
| Pelargonidin | 50-100 | 79 | 8.1 | 83 | 130 |  |
| Quercetin | 83.4 | 62 | 3.3 | 61 | 108 |  |
| Suramin | 20 | N.D | 1.2 | -4.8 | 91 |  |
| TGG | 2.86 | 12 | 13 | 51 | 80 |  |

<sup>a</sup>Literature reported EC<sub>50</sub> values performed in a variety of SARS-CoV-2 infectivity assays; see citations in **Table S1**. <sup>b</sup>Critical aggregation concentration determined by DLS. <sup>c</sup>IC<sub>50</sub> against counter-screen enzyme(MDH or AmpC\*). <sup>d-e</sup>Single -point Triton X-0.01%<sup>d</sup> and BSA (10mg/mL)<sup>e</sup> reversal assay performed at approximately MDH or AmpC\* IC<sub>90</sub>. All values reflect testing in triplicate, and errors of +/- 30%.

To be convincingly categorized as a colloidal aggregator, we asked if the molecules that formed particles by DLS also inhibited widely used counter-screening enzymes in a detergent dependent manner (i.e., inhibited in the absence of non-ionic detergent and had that inhibition much attenuated in the presence of detergent). All 41 drug-library compounds were thus tested in enzyme assays against malate dehydrogenase, initially at either 100 or 200 µM, depending on their solubility and their literature-reported activity (**Figure S3**). If ≥40% enzyme inhibition was observed, inhibition was measured in full concentration-response to determine IC_50_ values (**Figure 2**, **Table 1**). If compound did not inhibit MDH but formed colloidal-like particles by DLS, it was also assayed against AmpC (**Figure 2, Figure S3**). To confirm an aggregation-based mechanism, we investigated whether inhibition could be diminished by addition of 0.01% v/v Triton X-100 and by 10 mg/mL BSA, both of which disrupt colloid-based inhibition with little effect on well-behaved inhibitors [26, 63]. Of the 33 drug-library molecules that formed particles by DLS, 17 inhibited the counter-screening enzymes in a detergent- and BSA-dependent way, with inhibition being either fully or substantially reversed in the presence of these agents (**Table 1**). An exception was suramin, which did not form particles by DLS (**Figure S2**), did inhibit the counter-screening enzymes, was not detergent-dependent, but was sensitive to BSA (**Table 1)**; suramin appears to be a promiscuous inhibitor, but not aggregation based. Meanwhile, oroxylin A and EGCG also did not form colloidal particle by DLS (**Figure S2**), but did inhibit the counter-screening enzymes in a detergent-dependent fashion (**Table 1**); the status of these molecules as colloidal aggregators remains uncertain. In summary, 17 of the 41 drug-library molecules found to inhibit viral entry formed colloid-like particles at concentrations relevant to their literature reported activity, and inhibited counter-screening enzymes in a detergent- and a BSA-dependent manner, consistent with their status as colloidal aggregators.

**Figure 2.**
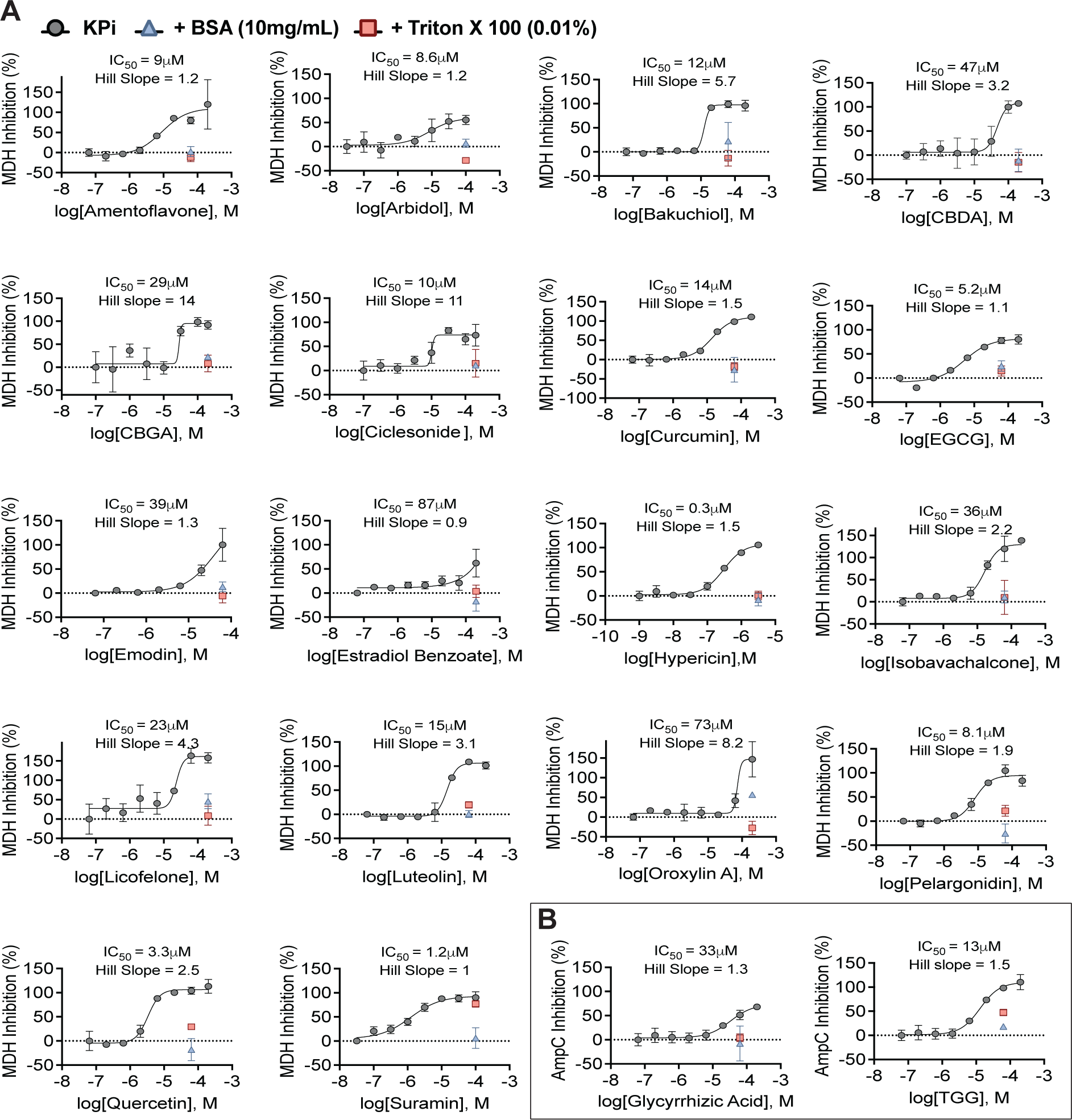
Detergent-dependent inhibition of the counter-screening enzymes (**A**) MDH or (**B**) AmpC. Percent inhibition after pre-incubation with BSA (blue triangle) or the addition of Triton X-100 (red square). All measurements performed in triplicate.

### Exploring BSA-dependent attenuation on SARS-CoV-2 entry inhibitors

If BSA reverses colloid-based inhibition by the repurposed drug-library molecules, it might also be expected to reverse their inhibition of viral entry (we were unable to test detergent, which also reverses colloid-based enzyme inhibition, because it destabilized the viral particles on its own (**Figure S4**)). We tested 20 of the drug-library molecules for their ability to inhibit entry of a SARS-CoV-2 Spike pseudotyped lentivirus with a luciferase reporter gene in A549-hACE2-TMPRSS2 cells; this pseudovirus has been widely used to mimic the behavior of SARS-CoV-2 without using a fully infectious form of the virus [64]. Sixteen of the colloidal aggregators tested showed loss of infection on co-treatment with BSA at a single concentration (**Figure 3A**). For six of these, we also looked at the effect of BSA in concentration-response. All six suffered at least a 10-fold drop in potency, with most displaying even greater potency losses such that no residual antiviral efficacy was measurable in the presence of BSA (**Figure 3B**). We note that EGCG, which behaved as a colloidal aggregator in the detergent- and BSA-dependent counter-screens, but did not form detectable particles by DLS, also showed no inhibition against viral entry upon the addition of BSA (**Figure 3A-B**). Consistent with the aggregation-bases of entry inhibition of the drug-library molecules, a well-studied colloidal aggregator, sorafenib, which may be considered a positive-control, also inhibited viral entry with an EC50 of 0.41 µM, a value that increased 50-fold in the presence of BSA (**Figure 4C**). Conversely, the inhibition of viral entry by what is effectively a negative control molecule, azelastine, which is not a colloidal aggregator, was little affected by the addition of BSA (**Figure 4**).

**Figure 3.**
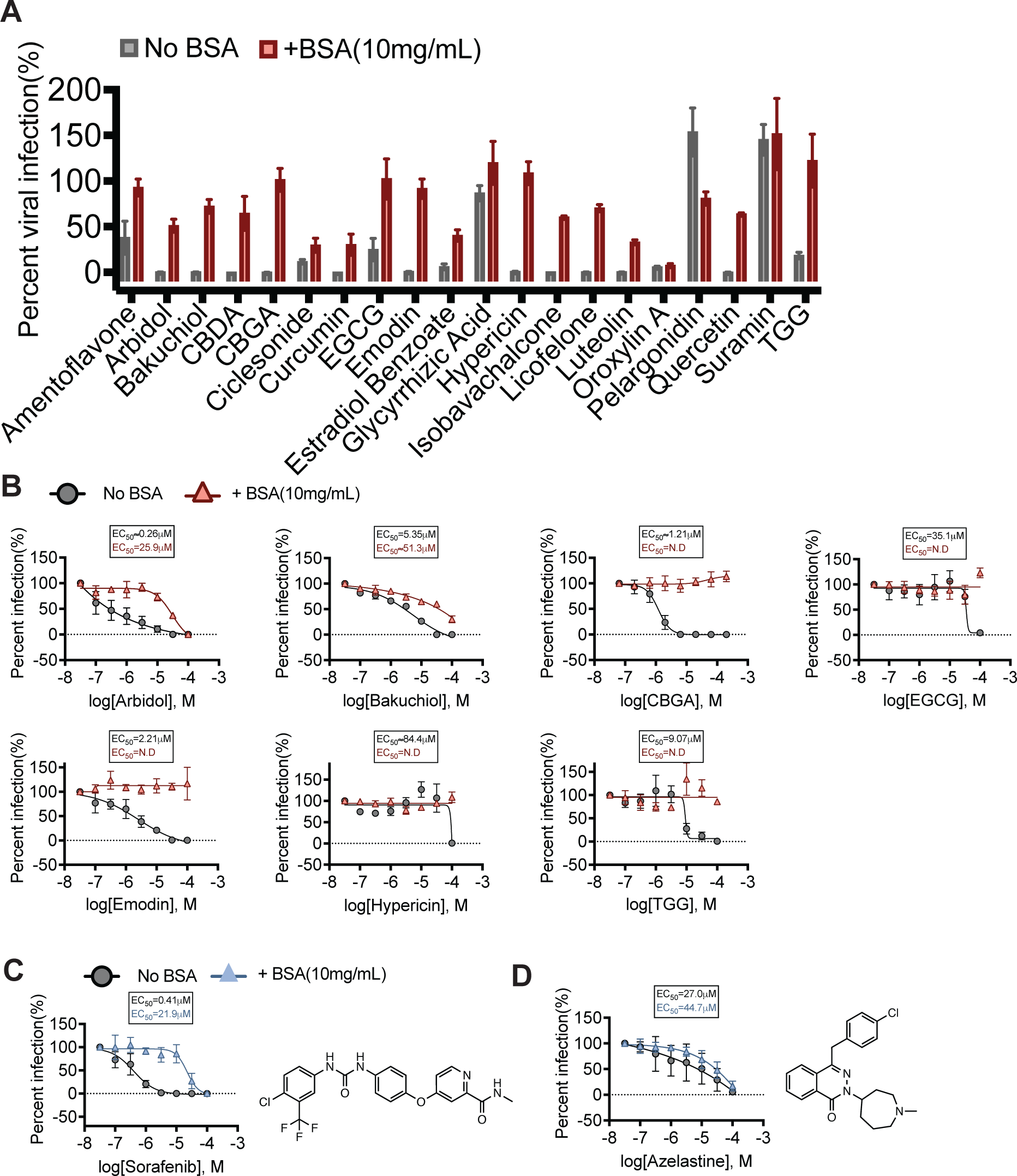
Spike Pseudovirus infectivity assay with BSA perturbation. (**A**) Percentage of viral infection with treatment of identified colloidal aggregator hits tested at a single-point concentration (63.3µM or 31.6µM) in the presence of BSA (red) or without (gray). (**B**) Percentage of viral infection with treatment of colloidal aggregator and nonspecific inhibitor(EGCG) in the presence of BSA (red) or without (gray). (**C**) Known aggregator and (**D**) Non-aggregator in the presence of BSA (blue) or without (gray). All data shown was performed in A549-hACE2-TMPRSS2 cells for three independent experiments performed in triplicate.

**Figure 4.**
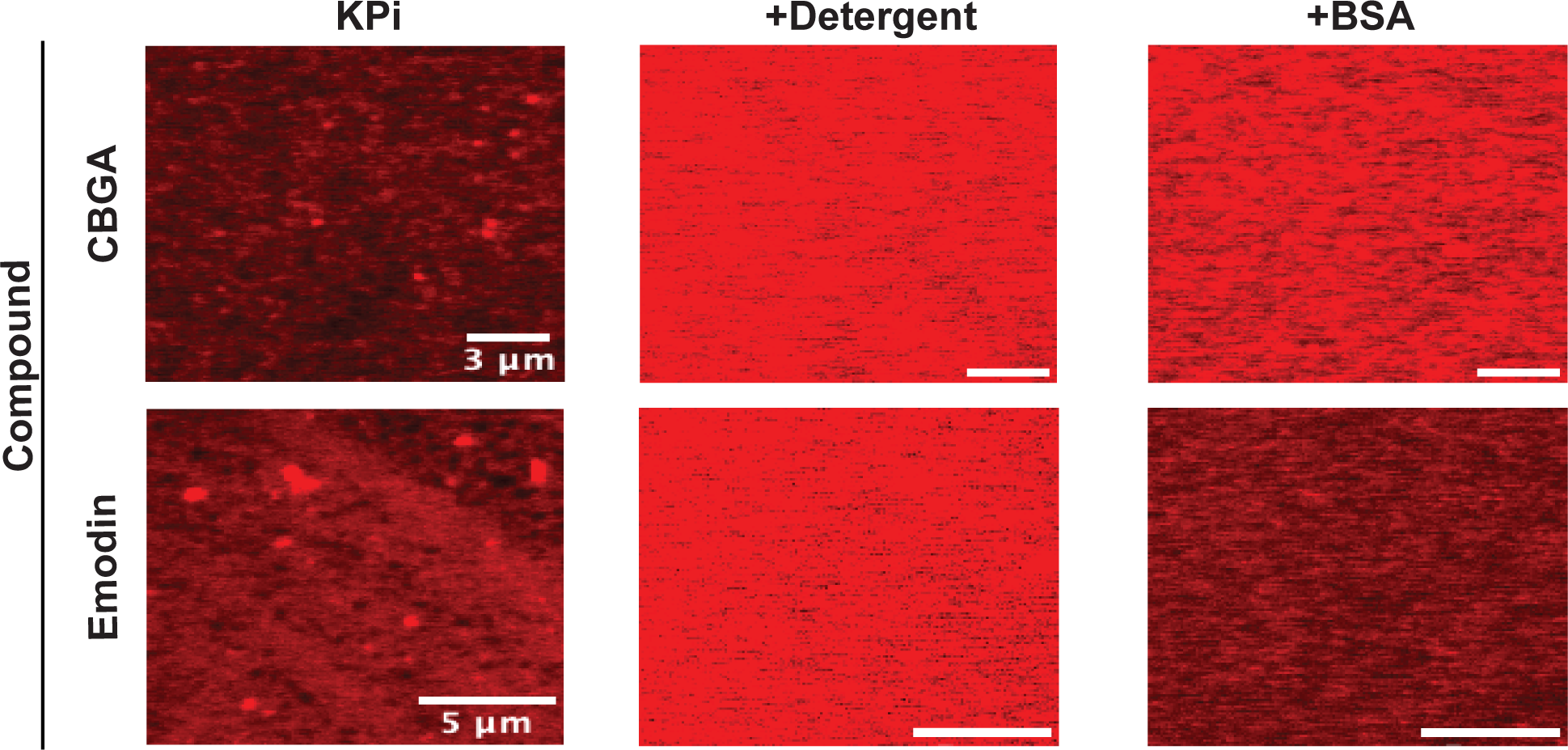
Spike protein binds to colloidal aggregates. Spike protein labeled with Alex-fluor-647 forms punctate fluorescence in the presence of the colloidal aggregators CBGA and emodin by confocal fluorescence microscopy, consistent with the viral protein binding to the colloidal particles. In the presence of the non-ionic detergent Tween-80 (0.025%) or of 10 mg/mL BSA, the puncta disappear, and the images return to the disperse fluorescent characteristic of fluorescent spike protein alone in phosphate buffer. Colloidal aggregates visualized with Spike protein alone in KPi and in the presence of detergent (0.025% Tween-80) or BSA (10mg/mL).

### Imaging colloid protein sequestration with fluorescently labeled Spike protein

Viral entry inhibitors are thought to work by binding to the spike protein of SARS-CoV-2. Colloidal aggregators might be expected do the same, though here one would expect it would be many thousands of spike proteins binding to each individual colloidal particle, as has been shown with other proteins [31, 36]. Using Spike protein fluorescently labeled with Alexa-fluor 647 we sought to visualize such protein-colloid complexes directly. Labeled spike protein alone displays as a disperse field of red fluorescence, as expected. On addition of two of the colloidal aggregators identified in this study, emodin and CBGA, the spike protein fluorescence became punctate, consistent with sequestration on the colloidal drug particles (**Figure 4**), and as previously seen for green fluorescent protein binding to well-established colloidal aggregators [26]. On addition of detergent, which disrupts colloidal particles, the fluorescent puncta disappeared. Similarly, pre-incubation of the colloidal drug particles with BSA, which prophylactically coats aggregates, prevented puncta formation (**Figure 4**). Finally, no puncta were observed in the presence of the non-aggregator azelastine (**Figure S5**). These observations are consistent with direct binding of the spike protein of SARS-CoV-2 to colloidal aggregates of molecules from the drug library, and suggests a mechanism for their action.

## Discussion

From this study of the role of colloidal aggregation in antiviral cell-based screens, three key observations emerge. **First**, colloidal aggregators are common in these screens, with 17 out of the 41 drug-library compounds investigated meeting all of our criteria for classic aggregators: formation of 50 to 1000 nm particles by DLS, inhibition of counter-screening enzymes, and reversal of this inhibition by agents like 0.01% Triton X-100 and BSA, and often high Hill slopes in the inhibition curves. As in other examples of aggregation-based inhibition, the mechanism appears to be sequestration of the active protein, here likely spike protein, on the colloid surface (**Figure 4**). **Second**, colloidal aggregation can confound in cell-based assays. Small molecule aggregation has typically been observed and controlled for in pure protein assays, it is less commonly looked for in cell-based assays [65]. Here, aggregation may be sequestering the spike protein of SARS-CoV-2 (**Figure 4**), and so inhibiting overall viral entry, but this confound may occur more broadly in cell-based assays. **Third** and more hopefully, this study suggests rapid controls for colloidal aggregation in cell-based assays. While the addition of detergent is a simple way to check for aggregation-based activity in pure protein assays, cellular assays do not always tolerate it, and in anti-viral assays even non-ionic detergent can disrupt the viral particles themselves, making detergent problematic as a control. Fortunately, BSA can also be used if not to disrupt colloidal aggregates, then to prophylactically coat them, preventing the association of viral or other target proteins, and so saving them from artifactual effects.

Certain caveats should be mentioned. The compounds tested were selected with some bias—we looked for molecules that resembled aggregators among the reported repurposed drugs—and we do not claim the hit rate achieved here will translate to unbiased controls of viral entry inhibitors. Nor do not believe that colloidal aggregation is a concern in every context. For example, many assays are performed in the presence of 10% fetal bovine serum (FBS), which contains sufficient serum proteins to pre-saturate most colloids. Finally, BSA as a control reagent can have its own liabilities, as it is known to bind other small molecules via classical mechanisms, and so might perturb the apparent inhibition of even well-behaved molecules.

These caveats should not obscure the main observations of this study. Many of the drugs repurposed for Covid-19 in cell-based screens suffer from the artifact of colloidal aggregation and have dubious prospects as antivirals. Although we focus here on Covid-19, many of the same molecules are reported as antivirals against other viruses [60, 66, 67], implicating aggregation as a source of false-positives in past and, without the proper controls [68, 69], future antiviral drug discovery. More encouragingly, controlling for this widespread artifact is straightforward, via counter-screens of control enzymes or, more directly, addition of serum albumin into the assay. Those molecules whose cell-based activities are much reduced by the addition of BSA should be considered with suspicion, as they are likely acting via one of the most widespread artifacts in early drug-discovery, colloidal aggregation.

## Experimental Section

### Literature search

To identify drug candidates to screen for colloidal aggregation, we searched the literature using the keywords: “SARS-CoV-2”, “entry inhibitors”, and “drug repurposing”. Identified compounds were active in the micromolar range in cell-based assays. Compounds were visually inspected for characteristics of other known aggregators (e.g., multiple conjugated rings and planar compounds), while compounds with clogP ≤ 1 were filtered out. If a compound was an already reported aggregator and met the criteria above, it was also added to the drug candidates list.

### Compounds

All compounds were supplied as >95% pure as reported by the vendors, and were used as supplied without further purification. Compounds were ordered from Sigma Aldrich, SelleckChem, Cayman Chemical, TargetMol, Biorbyt, or Medchem Express.

### Enzyme inhibition assays

AmpC β-lactamase was purified as previously described [70]. Samples were prepared in 50 mM KPi buffer, pH 7.0 with final DMSO concentration at 1% (v/v). Compounds were incubated with 2 nM AmpC β-lactamase (AmpC) or Malate dehydrogenase (MDH) (Sigma Aldrich, 442610) for 5 minutes. AmpC reactions were initiated by the addition of 50 μM CENTA chromogenic substrate (Sigma Aldrich, 219475). The change in absorbance was monitored at 405 nm for 80 sec. MDH reactions were initiated by the addition of 200 μM nicotinamide adenine dinucleotide (NADH) (Sigma Aldrich, 54839) and 200 μM oxaloacetic acid (Sigma Aldrich, 324427). The change in absorbance was monitored at 340 nm, also for 80 sec. Initial rates were divided by the DMSO control rate to determine percent enzyme activity (%). Each compound was initially screened at 200 μM or 100 μM in triplicate, based on solubility and reported EC_50_ values of each compound. If a compound exhibited ≥ 40% inhibition against counter-screen enzyme, a concentration-response curve was performed to determine its IC_50_. Compounds that did not inhibit MDH but formed colloidal-like particles by DLS were screened against AmpC. Data was analyzed using GraphPad Prism software version 9.1.1 (San Diego, CA).

For detergent and bovine serum albumin (BSA) reversibility experiments, each compound that showed ≥40% inhibition was rescreened against counter-screen enzymes near its IC_90_. In detergent reversibility experiments, compounds were rescreened in the presence of 0.01% (v/v) Triton X-100 in triplicate. Enzymatic reaction was performed as described above. For BSA reversibility experiments, compounds were incubated in KPi buffer containing 10mg/mL BSA for five minutes to pre-saturate colloids [63]. The counter-screen enzyme was then added and incubated for another five minutes. Enzymatic reaction was performed as described above.

### Dynamic Light Scattering

Samples were prepared in filtered 50 mM KPi buffer, pH 7.0 with final DMSO concentration at 1% (v/v). Colloidal particle formation was detected using DynaPro Plate Reader II (Wyatt Technologies). All compounds were screened in triplicate at 200μM or 100μM. Initial screening concentration was determined off of literature reported EC_50_s. If colloidal-like particles were detected, eight-point half-log dilutions of compounds were performed in triplicate. As previously [31], critical aggregation concentrations (CACs) were determined by splitting the inhibition curves into two sections based on aggregating (i.e. >10^6^ scattering intensity) and non-aggregating (i.e. <10^6^ scattering intensity) points and were fitted with separate nonlinear regression curves. The point of intersection of the two curves was determined using GraphPad Prism software version 9.1.1 (San Diego, CA).

### Cell lines

A549-hACE2-TMPRSS2 (Invivogen, a549-hace2tpsa) and 293T (ATCC, CRL-3216) were maintained in Dulbecco’s Modified Eagle’s Medium (DMEM, Quality Biological, 112-319-101) supplemented with 10% Fetal Bovine Serum (FBS, Caisson Labs, FBL01) and 1X Penicillin-Streptomycin (Sigma-Aldrich, P4333). All cells were grown at 37°C and 5% CO_2_.

### Spike (SARS-CoV-2) pseudotyped lentivirus generation

To generate Spike pseudovirus, 293T cells were seeded at a density of 1X10^7^ in 15cm^2^ dishes in DMEM supplemented with 10% FBS and 1X Penicillin-Streptomycin. The next day, cells were transfected with pHAGE2-LUC2-ZsGreen, psPAX2, and SARS-CoV-2 Spike plasmids using Lipofectamine 3000 (Thermo Fisher, L3000015). 48 to 72 hours post-transfection, virus was harvested by collecting supernatant and filtering it through 0.45µm PES sterile filter (Thermo Fisher, 2954545). Virus was concentrated by ultracentrifugation with a 20% sucrose cushion. Pellet was resuspended in DMEM containing 1X Penicillin-Streptomycin, aliquoted, and stored at -80°C. The p24 level of pseudovirus stock was determined using a p24 ELISA kit using the manufacturer’s manual (TakaraBio, 631476).

### SARS-CoV-2 pseudovirus-based antiviral assay

To set up antiviral assays, compound stocks were added to 96-well white opaque bottom plate containing 150µL of DMEM (supplemented with 1X Penicillin-Streptomycin and Polybene (10µg/mL) (Sigma Aldrich, TR-1003-G) with final DMSO concentration at 1% (v/v). For BSA reversibility experiments, 10mg/mL Bovine Serum Albumin (Sigma Aldrich, 05470) was additionally added to DMEM and incubated for 5 minutes. Pseudovirus (10ng final) was then added/mixed followed by 40µL of A549-hACE2-TMPRSS2 cells and incubated for 24 hours. For luciferase reported assay, cells were lysed in 1X Passive Lysis Buffer (Promega, E1531) followed by Luciferase Assay Reagent (Promega, E1501). The luciferase activity was measuring using CLARIOstar (BMG Labtech) according to the manufacturer’s instructions (Promega). Percent infectivity was calculated by dividing sample luminescence by DMSO control luminescence.

### Confocal Imaging Experiments

Alexa Fluor 647-labled Spike protein was diluted in 50mM KPi, pH 7 to 0.2mg/mL. Compound stock was added to Eppendorf tube (200µM and 2% DMSO (v/v) final), containing 50mM KPi +/- Tween 80 (0.025% (v/v) final) and 0.4µg of spike protein. Spike and compound were incubated in buffer for 5 minutes. For BSA perturbation, compound was added at same concentration to KPi buffer containing BSA (10mg/mL) and incubated for 5minutes followed by the addition of spike protein and an additional 5minute incubation. After incubations, Eppendorf content for each condition were transferred to glass coverslips. Images were acquired using Nikon Ti Microscope (Inverted) equipped with a Plan Apo VC 100x / 1.4 Oil objective. Fluorescently labeled spike was visualized using Evolve Delta EMCCD camera with a 640nm excitation and ET700/75m emission filter.

## Acknowledgments

BKS is supported by grants from the US National Institutes of Health U19AI171110 and R35GM122481. M.O. is grateful for support from the National Institutes of Health **(U19AI171110)**, the James B. Pendleton Charitable Trust, the Roddenberry Foundation, P. and E. Taft, and the Gladstone Institutes. M.O. is a Chan Zuckerberg Biohub – San Francisco Investigator. We thank Prof. Aashish Manglik for gifting us Alexa Fluor 647-labeled spike protein.

