## Supplemental information for "Colloidal aggregation confounds cell-based Covid-19 antiviral screens"

*Corresponding author

**Contents of SI:**

1. Table showing all compounds tested
2. DLS concentration-response for 16 literature compounds that formed colloidal-like particles
3. DLS scattering intensities of non-aggregators and nonspecific inhibitors
4. Single-point concentration screen against counter-screen enzymes
5. Detergent de-stabilizes pseudovirus alone
6. Confocal imaging of non-aggregator azelastine

**Table S1.**

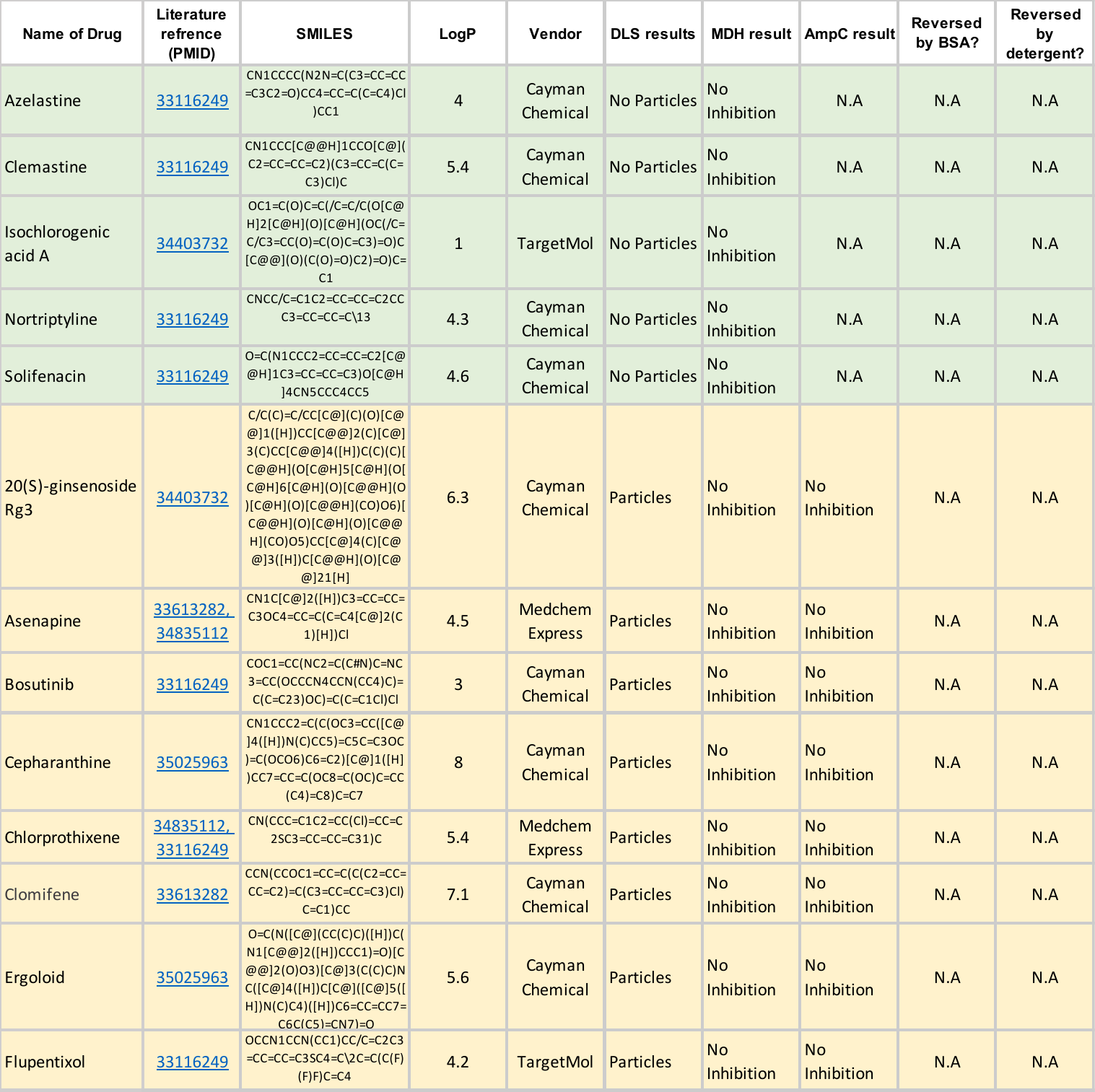
Drugs highlighted in red are considered aggregators, compounds highlighted in yellow satisfied some but not all criteria for aggregation, compounds highlighted in green are considered non-aggregators.

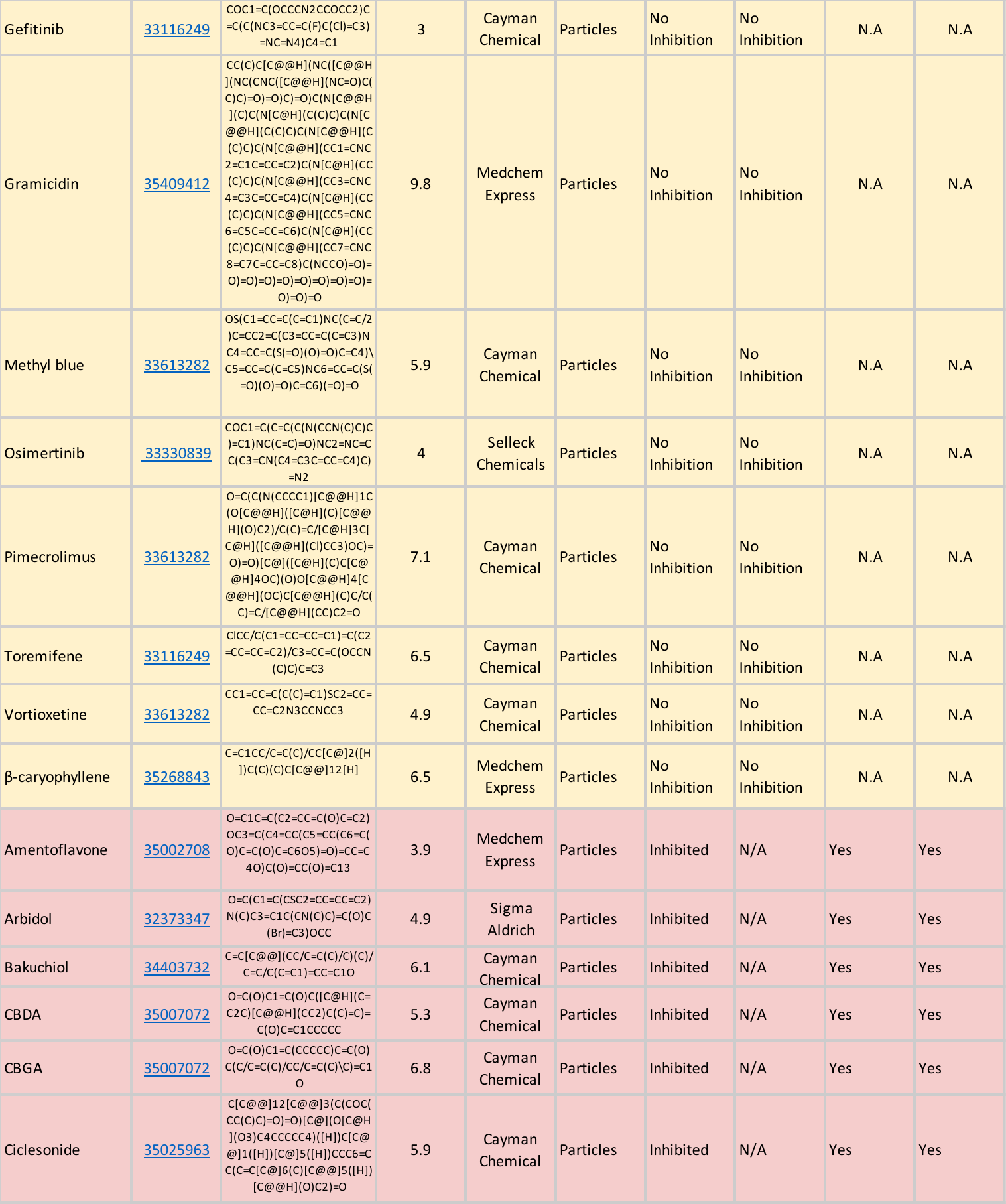

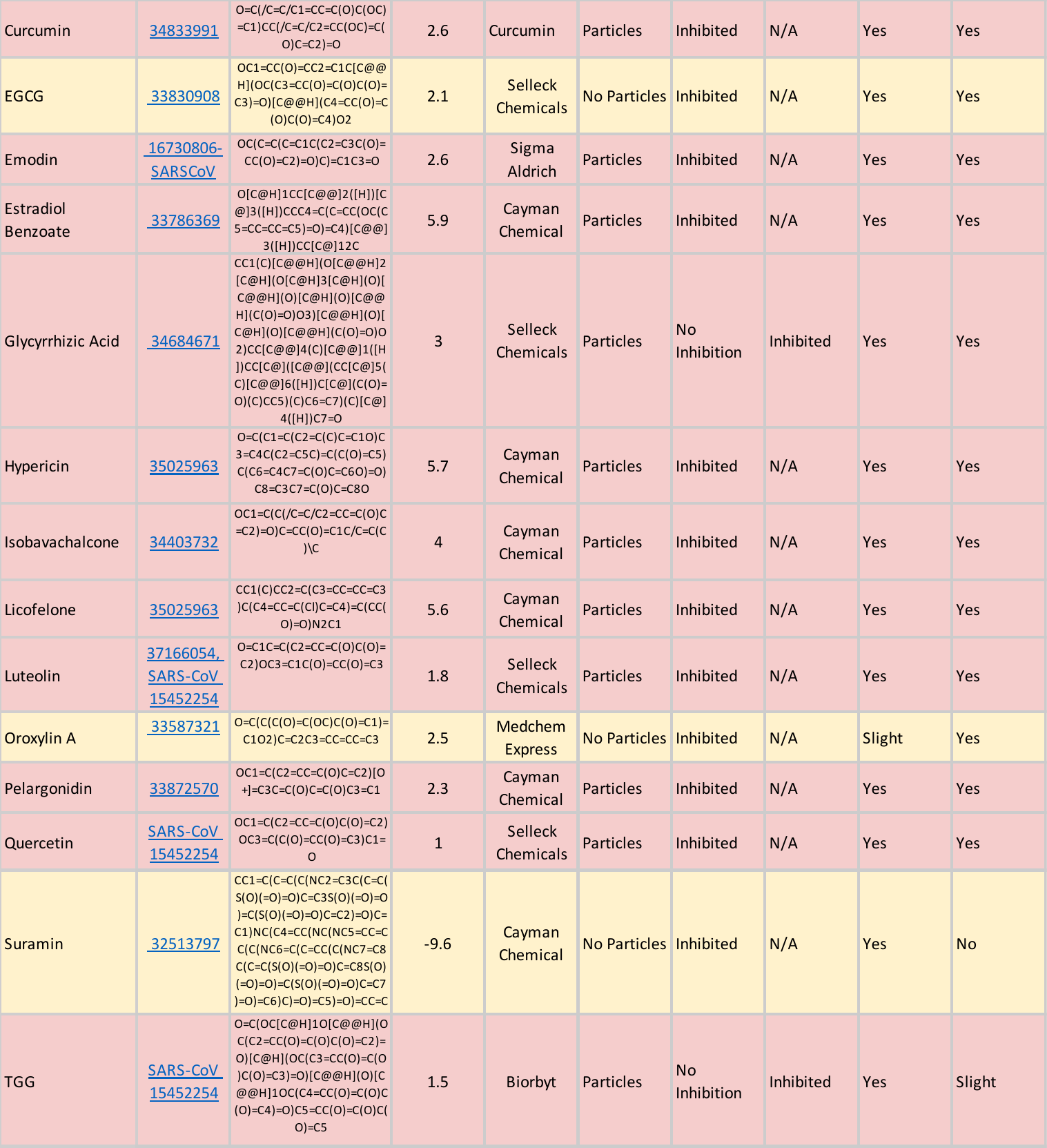

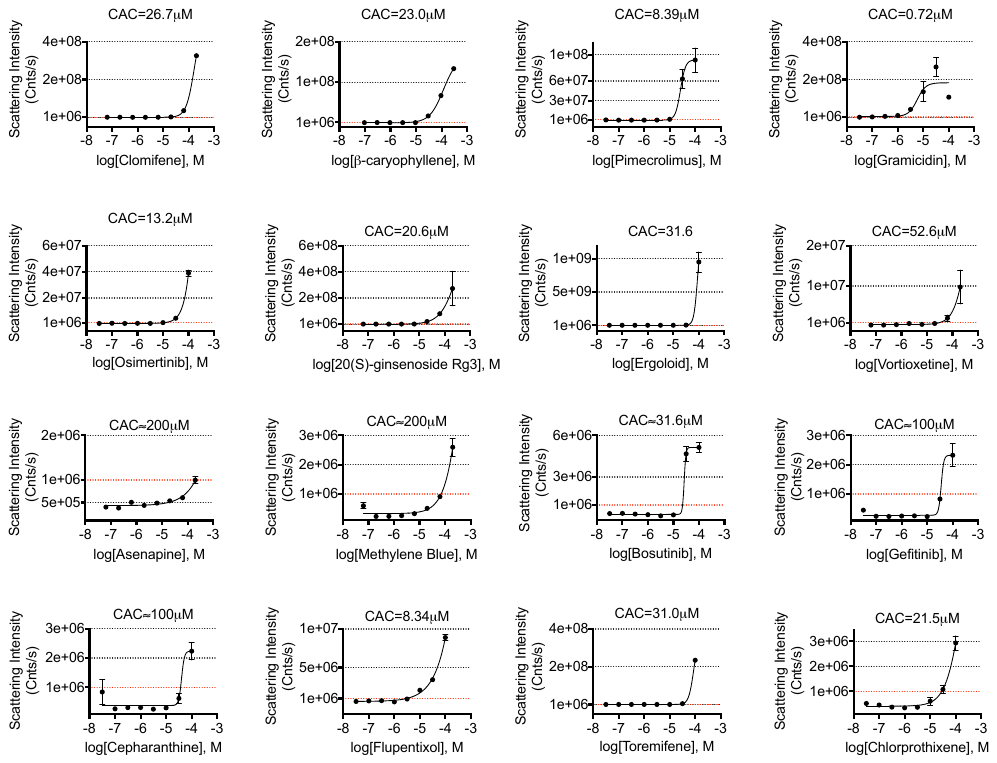

**Figure S1.** Dynamic light scattering performed on colloidal aggregator candidates that met scattering intensity threshold (1x10^6^) but did not inhibit counter-screen enzymes. Point of intersection serves as CAC value for each compound. All measurements were performed in triplicate.

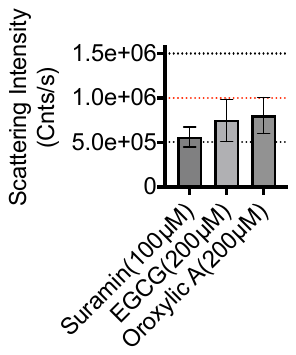

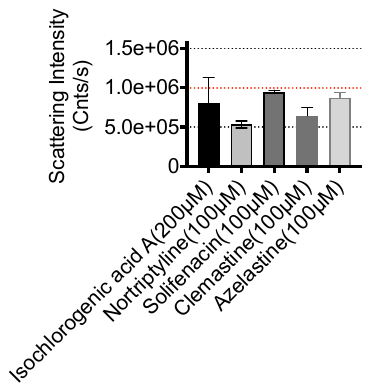

**B**

**A**

**Figure S2.** Dynamic light scattering of colloidal aggregator candidates that did not meet scattering intensity threshold of 1x10^6^. (**A**) Compounds that did not inhibit counter-screen enzymes. (**B**) Nonspecific inhibitors of MDH. All measurements were performed in triplicate

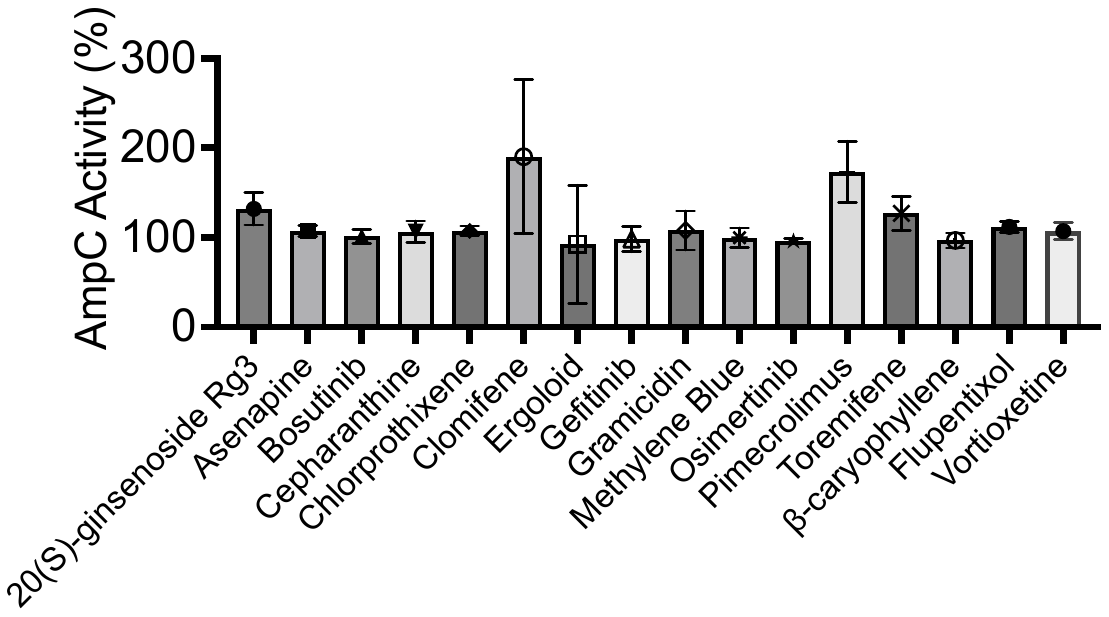

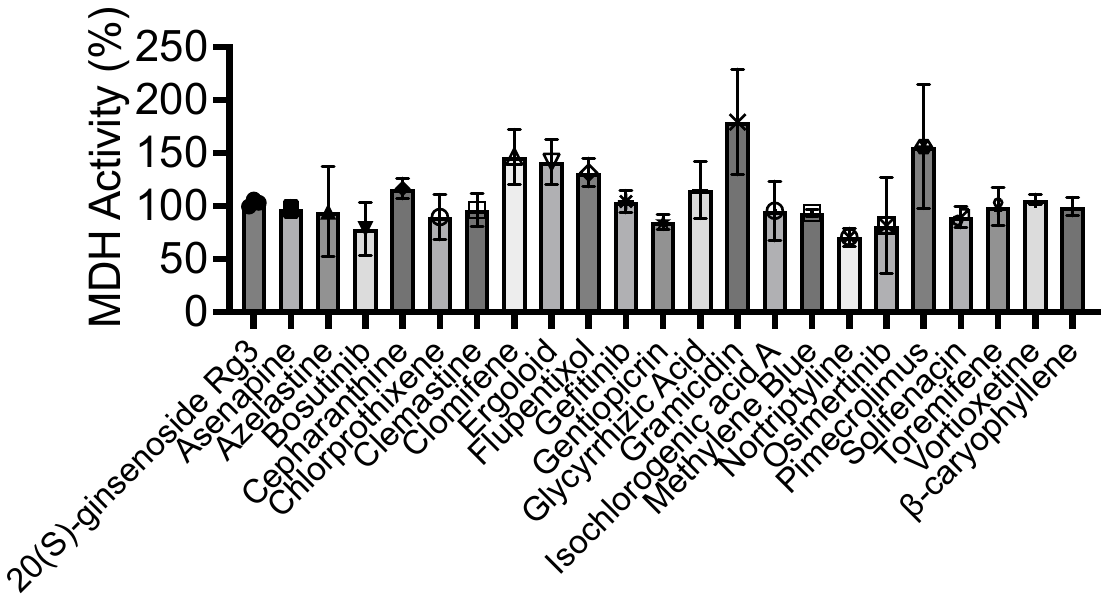

**B**

**A**

**Figure S3.** Enzyme inhibition assay against colloidal candidates. Single-point concentration of compounds (200µM or 100µM) screened against (**A**) MDH and (**B**) AmpC (if compounds formed particles by DLS). All measurements were performed in triplicate.

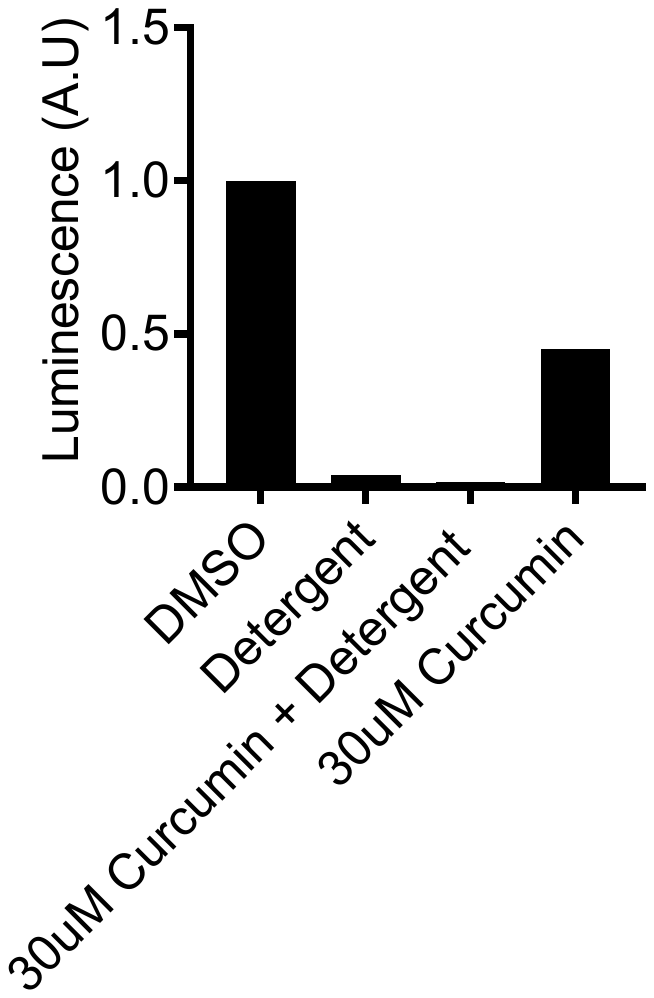

**Figure S4.** Spike Pseudovirus infectivity assay with detergent perturbation. Colloidal candidate, Curcumin, was screened at 30µM alone with pseudovirus and in the presence of Tween 80 (0.025% v/v). Tween 80 (0.025% v/v) and DMSO (1% v/v) was screened alone as controls.

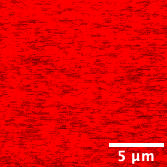

**Figure S5.** Visualization of spike protein and azelastine using confocal fluorescent microscopy. Spike protein labeled with Alex-fluor-647 in the presence of 200µM of non-aggregator, azelastine.
